# Using a recently approved tumor mutational burden biomarker to stratify patients for immunotherapy may introduce a sex bias

**DOI:** 10.1101/2021.05.28.446208

**Authors:** Neelam Sinha, Sanju Sinha, Kuoyuan Cheng, Sanna Madan, Ayelet Erez, Bríd M. Ryan, Alejandro A. Schäffer, Kenneth Aldape, Eytan Ruppin

## Abstract

The U.S. Food and Drug Administration (FDA) recently approved the treatment with pembrolizumab, an immune checkpoint inhibitor (ICI) targeting PD1 (anti-PD1), for patients with advanced solid tumors with a high tumor mutational burden (TMB) (defined as TMB ≥10 mutations/Mb). However, following recent studies suggest that TMB levels and response to ICI treatment may differ between male and female melanoma patients, we investigated whether using this high-TMB threshold for selecting patients for anti-PD1 treatment may induce a sex-dependent bias. We analyzed a large ICI cohort of 1,286 patients across nine cancer types treated with anti-PD1/PDL1. We find that using this threshold would indeed result in an unwarranted sex bias in melanoma, successfully stratifying female but not male patients. While this threshold is currently not a regulatory prerequisite for ICI treatment in melanoma, it is important to raise awareness to this bias. Notably, no sex-dependent significant differences were observed in the response of melanoma patients to anti-CTLA4 therapies, different chemotherapies or combination therapies. Beyond melanoma, the high-TMB threshold additionally introduces a sex bias of considerable magnitude in glioblastoma and in patients with cancers of unknown origin, however, these results are not statistically significant. A power analysis shows that these biases may become significant with larger sample size, warranting further careful testing in larger cohorts.

## Main Text

Treatment with immune checkpoint inhibitors (ICI) have shown remarkable clinical response in many cancers. This response is, however, limited to ∼15-20% of patients, raising a need for reliable response biomarkers especially biomarkers that apply to many tumor types to achieve maximum clinical benefits [1]. A biomarker increasingly referenced in clinical use is the tumor mutation burden (TMB), which is a measure of the total number of mutations in the coding region of the genome [2, 3]. A prospective biomarker analysis of the basket trial KEYNOTE-158, in which 1,066 solid tumor patients across 10 cancer types were treated with pembrolizumab, demonstrated that oncology patients with high-TMB, defined as ≥10 mut/Mb on the FoundationOne CDx assay, showed a higher frequency of response to anti-PD1 treatment vs non-high-TMB (<10 mut/Mb). The FDA subsequently approved the TMB ≥ 10 mut/Mb as a biomarker for administering anti-PD-1 therapy for advanced solid tumors that have progressed from prior treatment [4]. However, recent studies have suggested that the TMB levels, strength of immune selection, and response to ICI treatment differ between male and female melanoma patients [5-7]. These sex differences motivated us to examine whether usage of the 10 mut/Mb threshold for both sexes could introduce an unwarranted sex bias when selecting patients for anti-PD1 treatment.

To study this question, we mined the largest publicly available dataset of ICI-treated patient’ responses with TMB and demographic information [3]. This dataset includes 1,286 patients across nine different cancer types treated with anti-PD1/PDL1, 99 patients treated with anti-CTLA4 and 255 treated with an anti-PD1 + anti-CTLA4 combination. Among the 130 melanoma patients available in this cohort, we first observe a higher median TMB in male vs. female melanoma patients (median TMB=11.81 vs 6.51, respectively, Wilcoxon rank sum test P<0.10, Figure 1A top group), in concordance with previous reports [5]. We next asked whether the difference in survival of patients with high vs. non-high TMB is dependent on the sex of the patient. We find that using the ≥10 mut/Mb threshold identifies female melanoma patients with markedly better overall survival (hazard ratio (HR)=0.19, P<0.03), but fails to do so for male patients (HR=0.94, P<0.85, Figure 1B top group). The HR observed in male patients is thus five times higher than female patients (P-interaction between sex & TMB via log-rank test <0.03, Figure 1B top group).

**Figure 1:**
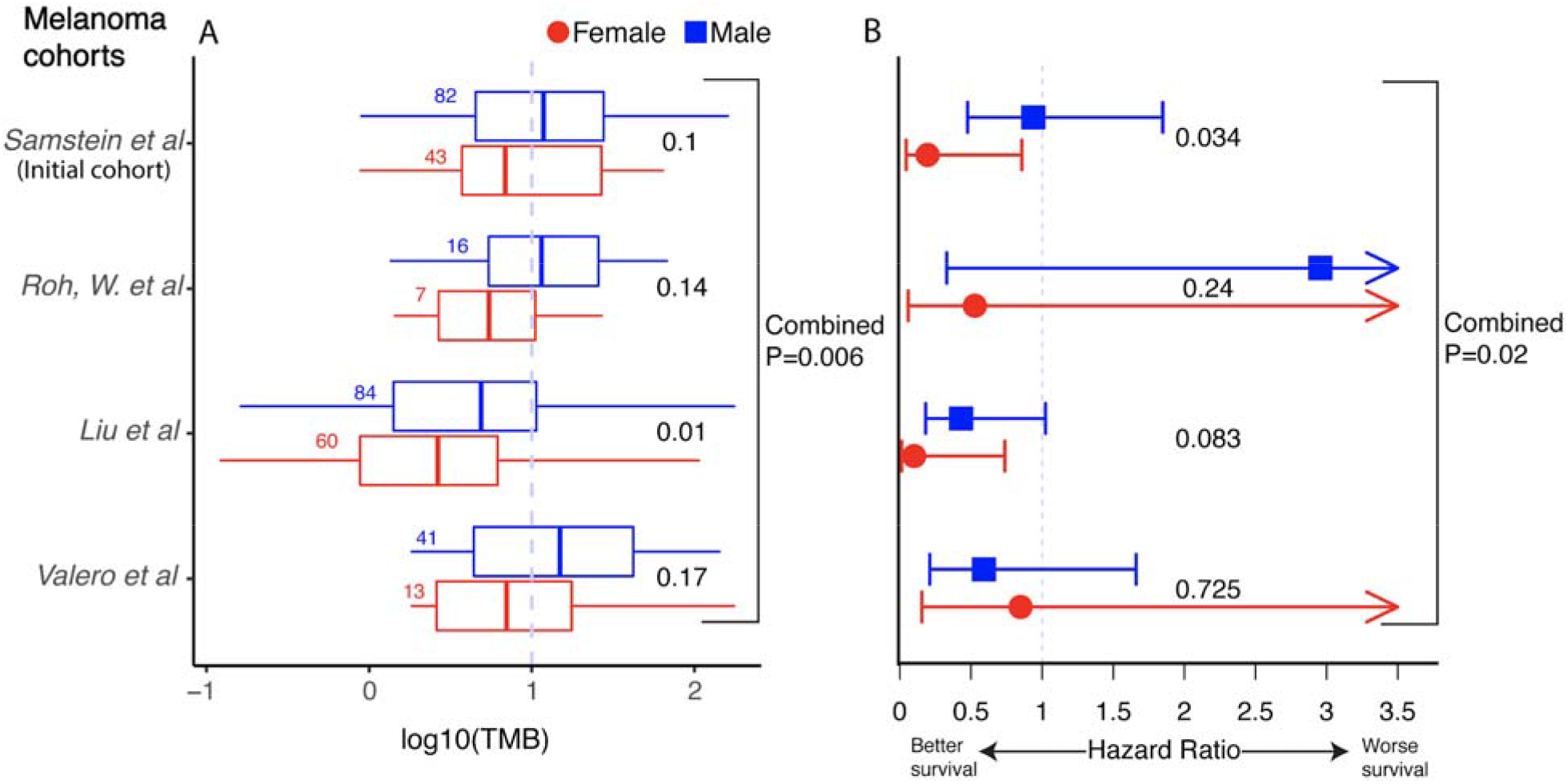
The association between high-TMB status and survival of melanoma patients after anti-PD1/PDL1 treatment is dependent on the sex of the patients. **(A)** The distribution of Log_10_(Tumor Mutation Burden (TMB)) the number of single nucleotide variants per Mb of sequenced genome (x-axis), for male and female patients for four different melanoma cohorts (Samstein et al., 2019, Roh et al., 2017, Liu et al., 2019 and Valero et al., 2021) (y-axis). The blue dotted vertical line denotes the FDA-approved TMB threshold for pembrolizumab of 10 mut/Mb. The number of samples in each group is provided alongside the respective box plots. The center line, box edges, and whiskers denote the median, interquartile range, and the rest of the distribution in respective order, additionally showing outliers. P-values of TMB differences are calculated using a one-tail Wilcoxon rank-sum test and provided on the right-hand side of each box plot. (B) Hazard ratios (HRs) for male (blue) and female (red) patients with high TMB (≥10 mut/Mb) vs the rest (x-axis) in four different melanoma cohorts (y-axis). Bars represent the standard 95% confidence intervals. The significance of difference in male vs. female hazard ratios is computed using a Wald test for the contribution of the coefficient of the interaction between TMB threshold and sex in a Cox proportional-hazards model.

To test the robustness of these findings, we repeated the above analysis in all additional publicly available melanoma cohorts treated with anti-PD1 where overall survival, TMB and patient demographics are available (Roh, et al., 2017 [8] (N=23), Liu et al., 2019 [9] (N=144) and Valero et al., 2021 [10] (N= 56)). Consistently, we observed a higher median TMB in male vs. female melanoma patients in each of these three cohorts (Figure 1A bottom three groups) and found a lower HR in female than male patients in two out of three cohorts, (Figure 1B bottom three groups). A combined meta-analysis (Weighted z-test) of all the four cohorts together shows a higher median TMB in male vs female patients (combined P=0.006) and a lower HR in female vs male patients (combined P=0.027). We note that these findings have limited immediate clinical implications as high TMB is not currently an FDA prerequisite for treating metastatic melanoma patients with anti-PD1 [11]. However, as clinicians may still take this threshold into account while considering therapies for a patient given the central role of TMB as a biomarker in general (and in ongoing clinical trials, e.g., NCT04187833, NCT02553642), we think it is important to take note of this potential bias.

We next tested whether the sex bias observed above extends to other ICI and non-ICI treatments in melanoma. To this end, we mined melanoma patients’ survival and TMB information in three additional patient cohorts, the first treated with anti-CTLA4 (N=174 [12, 13]), the second treated with an anti-PD1/PDL1 + anti-CTLA4 combination (N=115 [3]), and the third treated with different chemotherapies (N=322 [14]). We did not observe a significant difference in HR between male and female patients in any of these cohorts (P<0.14, P<0.8, P<0.4, in respective order), indicating that the sex bias observed in melanoma is specific to anti-PD1/PDL1 treatments.

We next asked whether the sex bias is present in other cancer types treated with anti-PD1/PDL1. Analyzing patient data across additional seven different cancer types from Samstein et al., 2019, we first charted the distribution of TMB values in tumors from female and male patients in each of these cancer types (Figure 2A). We observed considerable differences in the HR values between female vs male patients in glioblastoma (N=114, females vs. males HR=0.50 vs 0.89, P-Interaction<0.59, Figure 2B) and in cancers of unknown origin (N=88, females vs. males HR=1.03 vs 0.15, P-Interaction <0.06, Figure 2B). Notably, the HR is higher for males in glioblastoma patients and for females in cancer of unknown patients. The effect found in glioblastoma remained consistent when merging two additional small glioblastoma cohorts treated with anti-PD-1 (Zhao et al., ; N=15, Lombardi et al., ; N=12) [15,16] with our initial cohort (N=141, females vs. males HR=0.56 vs 1.19, P-Interaction<0.36).

**Figure 2:**
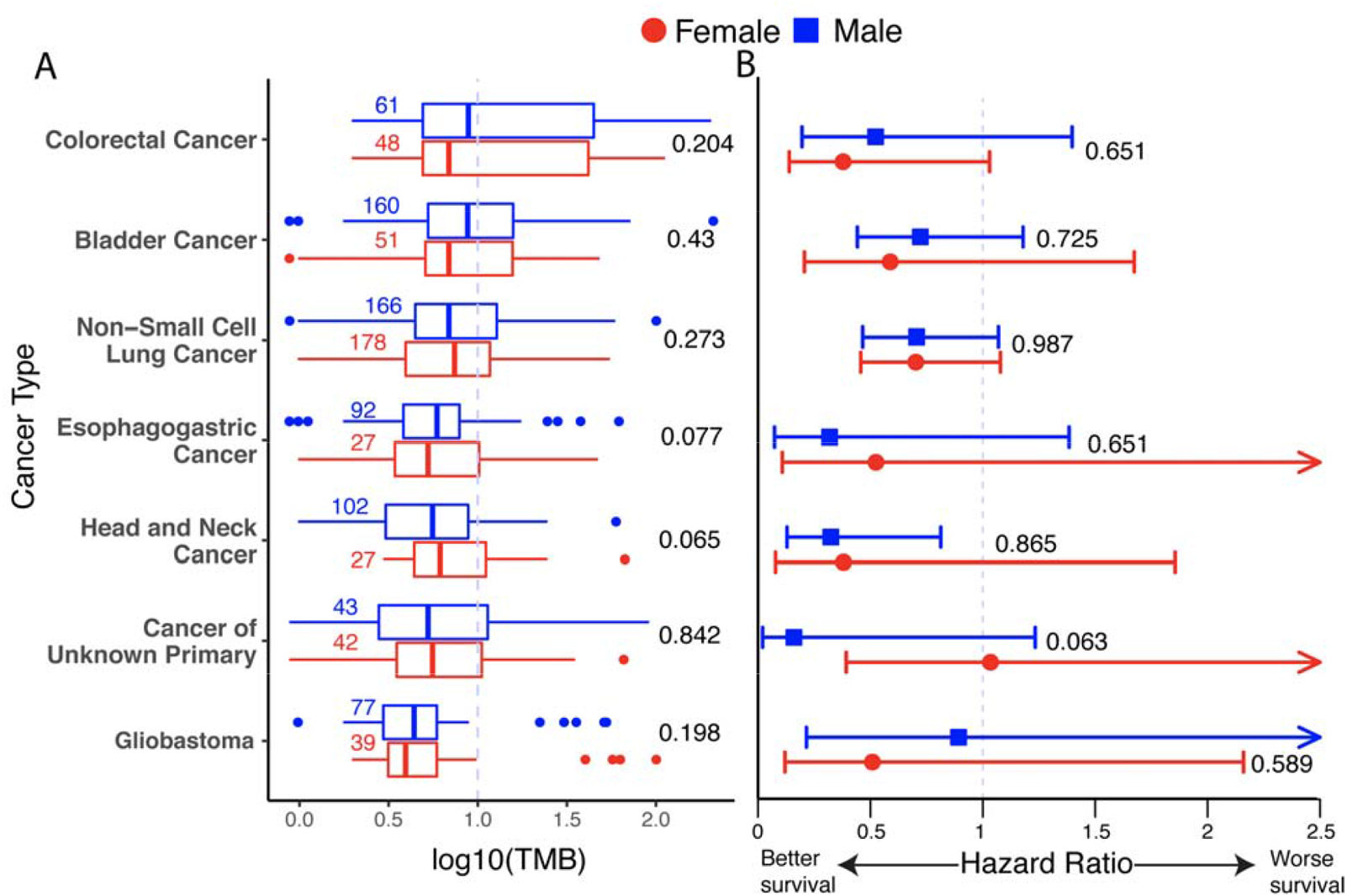
The association between high-TMB status and survival after anti-PD1/PDL1 treatment for male and female patients separately in seven cancer types. **(A)** Standard box plots displaying the distribution of Log_10_(TMB) (x-axis) for male and female patients across cancer types (y-axis), in a similar manner to Figure 1A. **(B)** Hazard ratios (HRs) of patients with high-TMB (≥10 mut/Mb) vs the rest (x-axis) in each cancer type (y-axis), sex color code as in [A]), displayed in a similar manner to that of Figure 1B. Renal cell carcinoma is not reported in our analysis as its HR cannot be computed confidently.

To test whether the small sizes of the glioblastoma and cancer of unknown origin datasets may impede the discovery of potentially significant sex-dependent effects, we down-sampled the melanoma anti-PD1/PD-L1 treatment cohort to the size of the glioblastoma and cancer of unknown origin cohorts (N=114 and N=88 respectively [3]). We repeated the down-sampling analysis 5,000 times, keeping the respective female to male ratio as in these cohorts. In these down-sampled melanoma cohorts, we find a large but statistically insignificant difference between HR in male and female patients: mean HR=0.20 and 0.95 for females and males, respectively; P-value=0.51 for a set size equal to that of glioblastoma cohort and a mean HR=0.20 and 1.04 for females and males, respectively; P-value=0.46 for a set size equal to that of cancer of unknown cohort. These results suggest that the small size of the glioblastoma and cancer of unknown origin may hinder our ability to identify significant trends and calls for further testing in larger cohorts.

Interestingly, we note that even though the size of the NSCLC cohort is substantial (N=329), we do not observe any notable difference of HR between male and female NSCLC patients (female vs male HR=0.70 vs 0.69, P-Interaction <0.99, Figure 2B), which is further confirmed in another cohort (N=16, interaction p<0.24) [17].

In summary, our findings indicate that the FDA-approved threshold of high-TMB for selecting patients for anti-PD1/PDL1 treatment is informative for stratifying female but not male metastatic melanoma patients. These findings may be of future relevance given ongoing clinical trials investigating the role of higher TMB as a biomarker for anti-PD1/PDL1 in melanoma (ClinicalTrials.gov Identifiers: NCT04187833, NCT02553642). Interestingly, in NSCLC, we did not observe any sex bias difference with TMB, despite the large size of the cohort. Further, our findings suggest that usage of this high-TMB biomarker may introduce a sex bias in glioblastoma and cancers of unknown origin, which needs to be carefully tested further in larger datasets as has been suggested by others for a variety of clinical findings regarding immunotherapy and immunology that may have a sex bias [18].

## Data and Code availability Statement

Scripts and data used in the study is provided to reproduce each step of results and plots in <u>this</u> GitHub repository.

## Acknowledgements

This research used the computational resources of the NIH HPC Biowulf cluster (http://hpc.nih.gov). This research was supported by the Intramural Research Program of the National Institutes of Health, NCI. S.S is supported by the NCI-UMD Partnership for Integrative Cancer Research Program.

## Author contributions

ER and SS conceived and supervised the study. NS and SS designed and developed the methodology. NS and SS acquired and analyzed the data. SS, NS, KC, SM, AAS, KS, AE, BMR and ER wrote the manuscript.

